# Inference of Multiple-wave Admixtures by Length Distribution of Ancestral Tracks

**DOI:** 10.1101/096560

**Authors:** Xumin Ni, Xiong Yang, Kai Yuan, Qidi Feng, Wei Guo, Zhiming Ma, Shuhua Xu

**Author notes:** Corresponding author (S.X.) and (Z.M.). These authors contributed equally to this work (Xumin Ni, Xiong Yang, Kai Yuan).

## Abstract

The ancestral tracks in admixed genomes are of valuable information for population history inference. A few methods have been developed to infer admixture history based on ancestral tracks. Nonetheless, these methods suffered the same flaw that only population admixture history under some specific models can be inferred. In addition, the inference of history might be biased or even unreliable if the specific model is deviated from the real situation. To address this problem, we firstly proposed a general discrete admixture model to describe the admixture history with multiple ancestral populations and multiple-wave admixtures. We next deduced the length distribution of ancestral tracks under the general discrete admixture model. We further developed a new method, *MultiWaver*, to explore the multiple-wave admixture histories. Our method could automatically determine an optimal admixture model based on the length distribution of ancestral tracks, and estimate the corresponding parameters under this optimal model. Specifically, we used a likelihood ratio test (LRT) to determine the number of admixture waves, and implemented an expectation??maximization (EM) algorithm to estimate parameters. We used simulation studies to validate the reliability and effectiveness of our method. Finally, good performance was observed when our method was applied to real datasets of African Americans, Mexicans, Uyghurs, and Hazaras.

## INTRODUCTION

Admixture among previously isolated populations has been a common phenomenon throughout the evolution of modern humans^1-3^. The history of population admixture has a strong influence on the landscape of genetic variation in individuals from admixed populations. Therefore, the population history of admixed populations can be reconstructed by utilizing genetic variation information^4-16^.

A few methods have been developed to infer admixture history based on ancestral tracks information^10-16^. Pool and Nielsen firstly used the length of ancestral tracks to infer population history^10^. They introduced a theoretical framework describing the length distribution of ancestral tracts and proposed a likelihood inference method to estimate parameters related to historical change in migration rates. Additionally, Pugach *et al.* performed wavelet transforms on the ancestral tracks in an admixed population to obtain the dominant frequency of ancestral tracks to estimate the admixture time^11,16^. Jin *et al.* further explored admixture dynamics by comparing the empirical and simulated distribution of ancestral tracks under 3 typical two-way admixtures models, i.e., the hybrid isolation (HI) model, gradual admixture (GA) model, and continuous gene flow (CGF) model^12^. They later deduced the theoretical distributions of ancestral tracks under HI and GA models^14^. Gravel extended these studies to multiple ancestral populations and discrete migrations, and provided a numerical estimation of tract length distribution^13^.

However, there was a significant shortcoming for all these methods. Before estimating the parameters of admixture history, a prior admixture model was required. The method by Pool and Nielsen considered a model that a target population received migrants from a source population^10^. Pugach *et al.*’s method was under an HI model, and Jin *et al.*’s methods were under HI, GA, and CGF models^11,12,14^. While Gravel considered models of multiple ancestral populations and discrete migrations, a prior admixture model was also required when dealing with the problem of admixture history inference^13^. However, in data analysis, we always have little information of admixture history, and the admixture model is often uncertain for some complex admixed populations^4,17-19^. Therefore, when the prior model deviates from the real history, these methods might be unreliable.

In our previous work^15^, we proposed some general principles in parameter estimation and model selection with the length distribution of ancestral tracks under a general model. However, with the increase of the number of parameters, it is complex and time-consuming to find the optimal solution, and too many parameters can lead to over-fitting. Thus, we only developed a method to infer admixture history under 3 typical two-way admixtures models.

In this work, we introduced a new method to select the optimal admixture model and estimate the corresponding parameters under a general model. Firstly, we proposed a general discrete admixture model with an arbitrary number of ancestral populations and arbitrary number of admixture events. This was similar to the general model in our previous work^15^. Then, we deduced the theoretical distribution of ancestral tracks with some reasonable approximations under the general discrete admixture model. We selected an optimal admixture model based on the length distribution of ancestral tracks. Specifically, we used a likelihood ratio test (LRT)^20,21^ to determine the number of admixture waves, and employed an exhaustion method to determine the order of admixtures. We then applied an expectation–maximization (EM) algorithm^22^ to estimate parameters under the optimal model. In our method, no prior knowledge about the admixture history was required, and the admixture model and its corresponding parameters could both be inferred by ancestral tracks. Finally, we conducted simulation studies to demonstrate the effectiveness of our method, and then applied our method to African Americans and Mexicans from the HapMap project phase III dataset^23^, and Uyghurs and Hazaras from the Human Genome Diversity Project (HGDP) dataset^1^.

## METHODS AND MATERIALS

### General Discrete Admixture Model

In our previous study^15^, we modeled admixture history generation by generation and proposed a general admixture model. The model was determined by a *K* × *T* admixture proportion matrix M = {*m_i_*(*t*)}_1≤*i*≤*K*,1≤*t*≤*T*_, where *K* is the number ofancestral populations, *T* is the time the admixed population arose, and *m_i_*(*t*) is the ancestry contribution of *ith* ancestral population at time *t*. If the admixed population did not receive any gene flows of *ith* ancestral population at time *t*, we set *m_i_*(*t*) as 0. This general model covers all scenarios of an admixed population with an arbitrary number of ancestral populations and arbitrary number of admixture events. However, the parameters for this general model are redundant, and will lead to over-fitting for most cases. For example, if we consider an HI model of 2 ancestral populations and the admixed population arose *T* generations ago, the number of parameters is 2*T*. However, 2(*T* – 1) parameters should be equal to 0, since ancestral populations contribute nothing after the first admixture. Thus, in fact, only 2 parameters must be estimated. Thus, to reduce redundancy and maintain the universality of our model, we proposed a general discrete admixture model that only records the information of actual admixture events (see Fig. 1).

**Figure 1.**
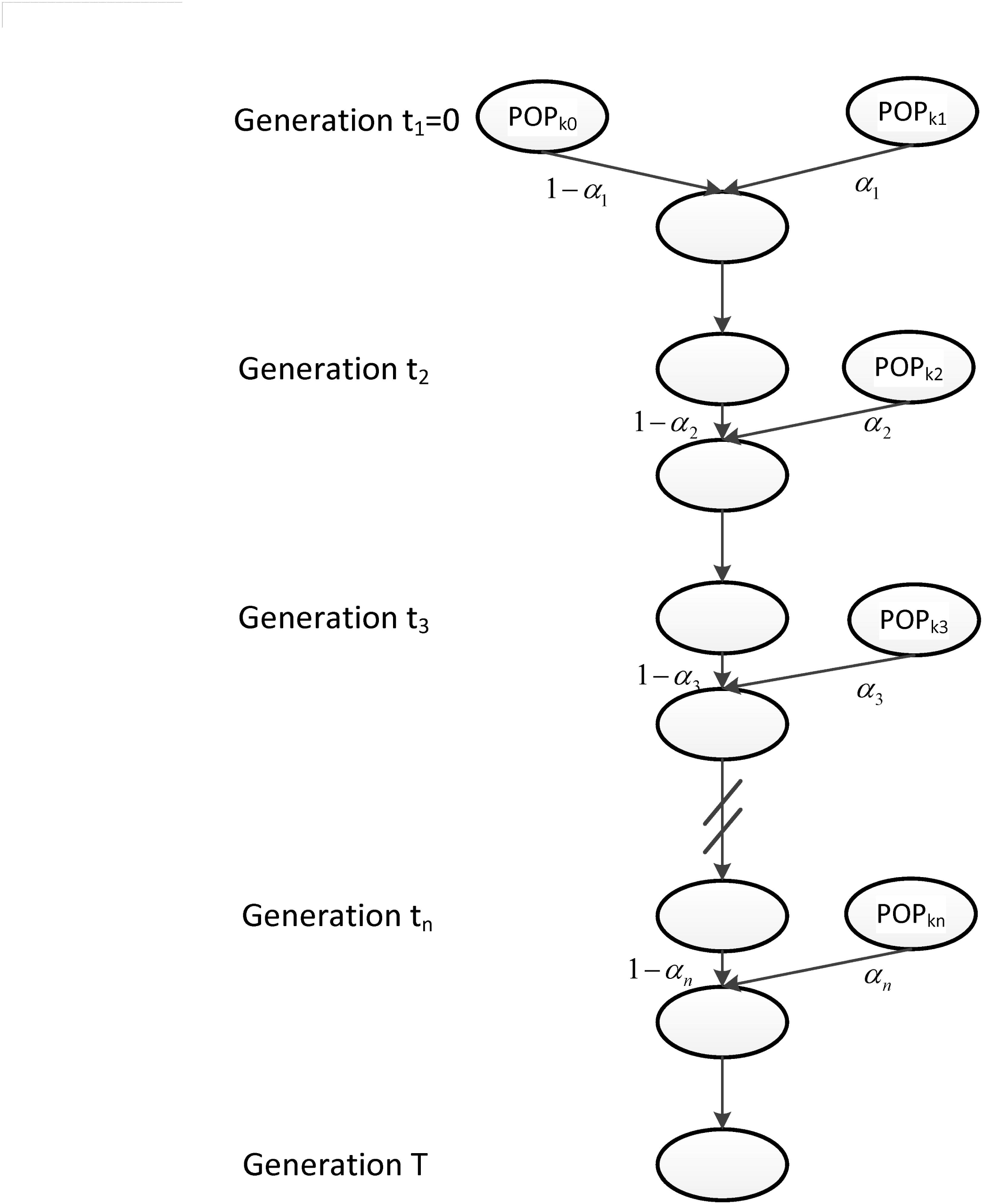
The general discrete admixture model. Here, we illustrated an admixed population with *K* ancestral populations and *n*-wave discrete admixtures, which started to admix *T* generations ago. POP_*ki*_ is the ancestral population of the *ith* admixture, is the admixture proportion of the admixture, and *t_i_* is the admixture time of the *ith* admixture.

We considered an admixed population with *K* ancestral populations and *n*-wave discrete admixtures. Here, the time of the admixture in generations increase over time, with *T* being the present time. For the first wave admixture (*i* = 1), there are 2 ancestral populations. We denote one ancestral population as population *k*_0_ and the other as population *k*_1_. When *i* ≥ 2, we denote *k_i_* as the ancestral population of *ith* admixture. Then, we denote a vector *0* = (*k*_0_, *k*_1_, …, *k_n_*) as the admixture order of ancestral populations. Let *α_i_* be the admixture proportion of the *ith* admixture and *t_i_* be the admixture time of the *ith* admixture. We note that 0 ≤ *α_i_* ≤ 1 for 1 ≤ *i* ≤ *n*, and *t*_1_ ≤ *t*_2_ ≤ … ≤ *t_n_* ≤ *t*_*n*+1_=: *T*. For convenience in our laterdescription, we denote the admixture event from population *k*_0_ as the 0*th* admixture, which means the ancestral population *k*_0_ is regarded as an admixed population before the first wave admixture. Thus, we set the corresponding admixture proportion *α*_0_ = 0 and admixture time *t*_0_ = *t*_1_. With this definition, each wave (*ith* wave) of admixture can be determined by 3 parameters, *k_i_*, *α_i_* and *t_i_*.

Now, denote *I_k_* = {*i*: *k_i_* = *k*}, then, *I_k_*(*j*) represents the wave ordinal of the *jth* admixture from ancestral population *k*. Let *n_k_* denote the number of admixture waves from ancestral population *k*, and thus we have *n_k_* = |*I_k_*| and 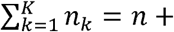 1, where *n* is total number of admixture waves. The general discrete admixture model is determined by the admixture order *O* = (*k*_0_, *k*_1_, …, *k_n_* the admixture proportion {*α_1_*}_0≤*i*≤*n*_, and the admixture time {*t*_0≤*i*≤*n*+1_. If we set

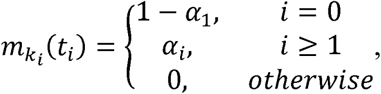

we can get the admixture proportion matrix of the previous general model^15^. This shows that our new model is similar to the previous model. Furthermore, this new model can also cover all scenarios of an admixed population with an arbitrary number of ancestral populations and arbitrary number of admixture events.

### Length Distribution of Ancestral Tracks

Next, we deduced the length distribution of ancestral tracks from ancestral population *k*. The deduction process is similar to that of our previous work^15^. The wave ordinals of admixture from ancestral population *k* are *I_k_*(1), *I_k_*(2), …, *I_k_*(*n_k_*), respectively. Denote *H_k_*(*t*) as the total ancestry proportion of the *kth* ancestral population in the admixed population at *t* generation, and then we have

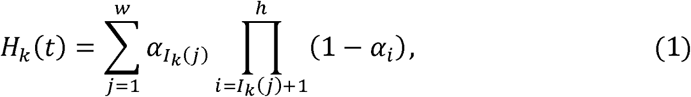

where *w* = max{*j*: *t*_*I_k_*(*j*)_ ≤ *t*, 1 ≤ *j* ≤ *n_k_* and *h* = max{*i*: *t_i_* ≤ *t*, 1 ≤ *i* ≤ *n*}.

Define *S_i_* as the survival proportion of the ancestral tracks from the *ith* admixture at generation *T*. Then,

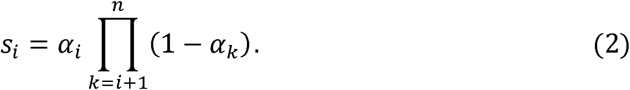

For simplicity, we assumed that chromosome length was infinite and there was no genetic drift. Additionally, we defined the recombination among tracks from different ancestral populations as effective recombination because we only observed these recombination events among different ancestries. The length of ancestral tracks was changed by these recombination events. For the tracks from ancestral population *k*, the effective recombination rate is 1 - *H_k_*(*t*) at *t* generation. Let *u_k_*(*j*) be thetotal effective recombination rate for ancestral tracks from the *jth* admixture of ancestral population *k*. Then, we have

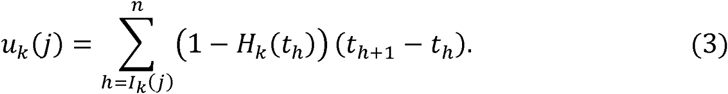

The length distribution of ancestral tracks from the admixture of ancestral population *k* is an exponential distribution with a rate of *u_k_*(*j*)^10,13,15^. A chromosome from the *jth* admixture of ancestral population *k* is expected to be split into *u_k_*(*j*) pieces per unit length (unit in Morgan). Thus, for the admixed population at *T* generation, the number of ancestral tracks from the *jth* admixture of ancestral population *k* is proportional to *s_I_k__*(*j*)*u_k_*(*j*). Let *X_k_* be the length of ancestral tracks from ancestral population *k* at generation *T*, and *f_k_*(*x*) is the probability density of *X_k_*. Then,

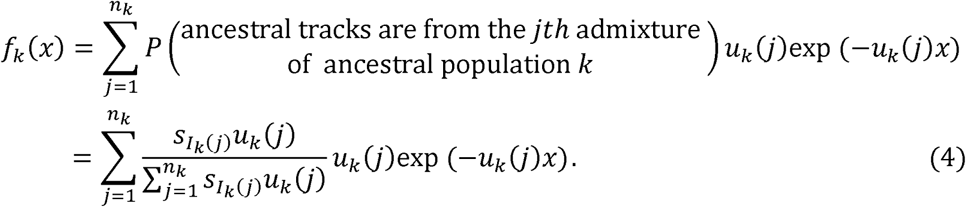

The length distribution of ancestral tracks was a mixed exponential distribution, and consisted with the results from our previous study^15^.

### Model Selection and Parameter Estimation

If the admixture model is determined, the length distribution of ancestral tracks can be written as Formula (4), which is a mixed exponential distribution. The EM-algorithm^22^ can be used to estimate the parameters in this distribution. However, the admixture model is often unclear in real situations, which means the number of admixture waves (*n_k_*) and the order of admixtures (*O*) are unknown. Thus, we must first determine *n_k_* and *O*. Here, we used LRT^20^ to select the optimal *n_k_.* After that, we used the exhaustion method to validate the accuracy of *O*. For any order of admixtures, we estimated the admixture proportion {aJosisn and admixture time {*α_i_*}_0≤*i*≤*n*_ and admixture time {*t_i_*}_0≤*i*≤*n*+1_ using the EM-algorithm. However, these parameter estimations must satisfy the following constraint conditions:

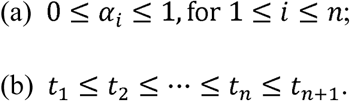

If the estimations don’t satisfy these conditions, the order is incorrect. After traversing all admixture orders, we could determine the correct ones.

The detailed procedures are as follows:

Step 1: Estimate the total admixture proportion *m_k_* of ancestral population *k*. With the inferred ancestral tracks, divide the total length of tracks from population *k* by the total length of tracks in the admixed population.
Step 2: Determine the number of admixture waves (*n_k_*) for each ancestral population and estimate the parameters of the mixed exponential distribution. For ancestral population *k*, use LRT to select the optimal number of admixture waves and then estimate the parameters of {(*ω*_*k*1_, *λ*_*k*1_), …, (*ω*_*kn_k_*_, *λ*_*kn_k_*_)} the mixed exponential distribution using the EM-algorithm, where 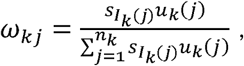 *λ_kj_*=*u_k_*(*j*). Details of the EM-algorithm and LRT procedures are in Supplementary Information (Supplementary Text S1).
Step 3: Select an admixture order *O* without replacement from set

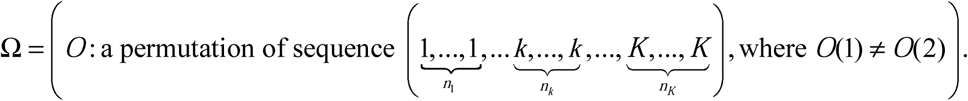 Get *I_k_* for each *k* base on the selected *O*.
Step 4: Determine {*S_i_*}_0≤*i*≤*n*_ and {*u_k_*(*j*), 1 ≤ *j* ≤ *n_k_*, 1 ≤ *k* ≤ *K*} from the following equations:

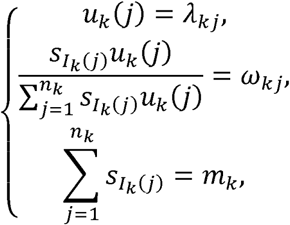

where 1 ≤ *j* ≤ *n_k_*, 1 ≤ *k* ≤ *K*.
Step 5: Determine {*α_i_*}_0≤*i*≤*n*_ from the following equations:

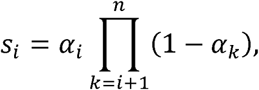

where 0 ≤ *i* ≤ *n*.
Step 6: Determine {*t_i_*}_0≤*i*≤*n*+1_ from the following equations:

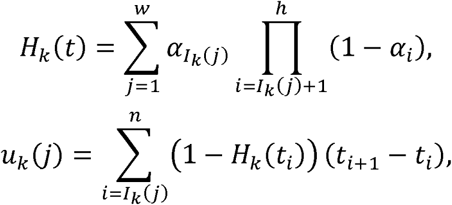

where 1 ≤ *j* ≤ *n_k_*, 1 ≤ *k* ≤ *K*.
Step 7: Judge whether {*α_i_*}_0≤*i*≤*n*_ and {*t_i_*}_0≤*i*≤*n*+1_ satisfy the following conditions:

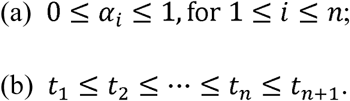

If these conditions are satisfied, record the corresponding admixture order *O*, admixture proportio {*α_i_*}_0≤*i*≤*n*_, and admixture time {*t_i_*}_0≤*i*≤*n*+1_. Then return to Step3 until all possible admixture orders are checked.

Through these above procedures, we obtained all reasonable admixture orders *O*, the corresponding estimators of admixture proportion {*α_i_*}_0≤*i*≤*n*_, and admixture time {*t_i_*}_0≤*i*≤*n*+1_. Based on the estimations of these parameters, we could recover the history of the admixed population.

However, due to a lack of accuracy in local ancestry inference, only these relatively long tracks are reliable^10,13^. Therefore, we are interested in the conditional length distribution of ancestral tracks longer than a specific threshold *C*. As we know, the length distribution of ancestral tracks from each ancestral population is a mixed exponential distribution. When we consider only tracks larger than *C*, the length distribution from ancestral population *k* becomes

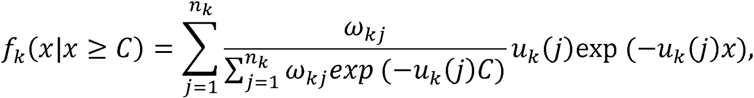

where 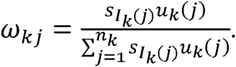 However, since this condition distribution is not a mixed exponential distribution, we cannot use the EM-algorithm to estimate the parameters. Fortunately, when we consider the random variable *Y_k_* = *X_k_* − *C*, we find that the distribution of *Y_k_* is a mixed exponential distribution, which can be written as follows:

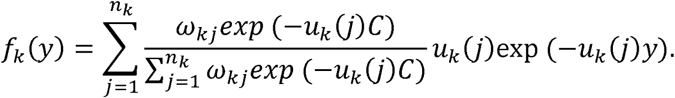

To take the threshold *C* into consideration, we must change the procedures of the aforementioned Step 2. We can easily obtain samples of *Y_k_* from samples of *X_k_*. Then, we can use the EM-algorithm and LRT to obtain the distribution parameters of *Y_k_*. Furthermore, by the relationship between *f_k_*(*x*) and *f_k_*(*y*), we can obtain the parameters of the mixed exponential distribution of *X_k_*. Then, the following Steps are the same as those aforementioned in Steps 3-7. These procedures were all implemented in our *MultiWaver*.

In the software of *MultiWaver*, 2 estimations of admixture time for the first wave were output. One was an estimation of *t*_0_, while the other was an estimation of *t*_1_. In theory, *t*_1_ is equal to *t*_0_, but in real data analysis, the estimations may be not equal because of random errors and tracks inference errors. Thus, we presented 2 estimations of admixture time for the first admixture wave in our results.

## SIMULATION

### Performance Evaluation of *MultiWaver*

We conducted simulations to evaluate the performance of *MultiWaver*. The simulation data were generated by forward-time simulator *AdmixSim*^24^. General settings of our simulation were the same as those in our previous study^15^.

Here, we divided multiple-wave admixture models into 2 different types of models. We denoted the model as a simple model if each ancestral population could contribute only once to the admixed population. The others were denoted as a complex model. In the complex model, at least 1 ancestral population donates more than once admixture. It is important to note that when we infer the admixture history under the complex model, it is very challenging to distinguish the different admixture waves from the same ancestral population.

We focused on evaluating the performance of *MultiWaver* under these 2 types of models. For the simple model, we considered a scenario of 3 ancestral populations (Fig. S1 Scenario (I)), and a scenario of 5 ancestral populations (Fig. S1 Scenario (II)). For the complex model, we considered a scenario of 2 ancestral populations with 2-wave admixtures (Fig. S1 Scenario (III)). We evaluated the performance of *MultiWaver* with different admixture times and admixture proportions. For simplicity, we supposed the admixture proportions (*α_i_*, 1 ≤ *i* ≤ *n*) were equal. We set 3 different values of admixture proportion, 0.1, 0.3, and 0.5, for each scenario. For Scenario (I), the admixture time was set as 2 different cases: (a) *t*_2_ = 20, *T* = 40, and (b) *t*_2_ = 40, *T* = 60. For Scenario (II), the admixture time was also set as 2 different cases: (a) *t*_2_ = 20, *t*_3_ = 40, *t*_4_ = 60, *T* = 80, and (b) *t*_2_ = 40, *t*_3_ = 80, *t*_4_ = 120, *T* = 140. However, for Scenario (III), the admixture time was set as 4 different cases: (a) *t*_2_ = 20, *T* = 40, (b) *t*_2_ = 40, *T* = 60, (c) *t*_2_ = 60, *T* = 80, and (d) *t*_2_ = 80, *T* = 100. Each case was repeated 10 times for a total of 240 simulations across these 3 scenarios. *MultiWaver* was applied to the simulated data with the default settings; the results were recorded and summarized.

In real situations, due to the limitations of local ancestry inference, only the ancestral tracks longer than a special threshold can be accurately inferred. Thus, to make our method more available to real situations, we chose the thresholds ranging from 0 cM to 2 cM in steps of 0.25 cM, and then evaluated the robustness of our method under different thresholds.

### Application to Real Datasets

Firstly, we applied our method to some real datasets of African Americans and Mexicans. These 2 populations are typical admixed populations and their histories are relatively clear. Therefore, they could be used to test the performance of our method for real data. We obtained the datasets of African Americans (ASW), Mexicans (MEX), and reference populations African (YRI) and European (CEU) from the HapMap Project Phase III dataset^23^. Meanwhile, Maya and Pima populations represented American Indian ancestry, which were obtained from the HGDP dataset^1^. According to prior knowledge, African Americans and Mexicans have more than 2 ancestries^14,25^. However, the proportion of Native American ancestry of African Americans is less than 5%^26^, and thus, we only considered 2 dominant ancestries (African and European ancestry) of African Americans. For Mexicans, we considered 3 ancestries: African, European, and American Indian ancestry^27^.

Then, our method was used to reconstruct the population history of Uyghurs and Hazaras. The histories of these 2 populations are more complex. Uyghurs and Hazaras populations were obtained from the HGDP dataset. Previous studies have shown that Uyghurs and Hazaras had admixed ancestries mainly from Europe and East Asia^1,5^. Here, we used Han and French as the proxies of Asian ancestry and European ancestry, respectively^8^. These reference populations were also obtained from the HGDP dataset.

To enhance the reliability of our analysis, HAPMIX^6^ was selected as the local ancestry inference method since it shows good performance in admixture break points inference^28^. However, HAPMIX can only be used to detect ancestral tracks for two-way admixtures, and thus it might not be proper for the Mexican population. PCAdmix^29^ has shown great power in inferring the local ancestry of Mexican populations^27^, and thus we used PCAdmix in this study. The generations pre-set in HAPMIX inference were 10 for African Americans and 80 for Uyghurs and Hazaras. The window size set in PCAdmix was default. Since phasing data was required for both HAPMIX and PCAdmix, SHAPEIT 2^30^ was used to infer the haplotype phase. Finally, *MultiWaver* was used to determine the optimal model and estimate the admixture time accordingly with tracks longer than 1 cM.

## RESULTS

### *MultiWaver* Performed Well under Simple and Complex Models

We compared the admixture histories inferred by *MultiWaver* with the histories set in simulations, and then evaluated the performance of our method in model selection and parameters estimation. For Scenario (I) and (II), results showed that estimations of admixture time were high consistency with the time simulated if we pre-set the admixture model as the simple model (-s option in *MultiWaver)* (see Fig. 2). Our method also performed well when we did not pre-set this option (-s) (see Fig. S2). Only a few models in our simulations were wrongly selected. When the model was correctly selected, the admixture time estimated was consistent with the simulated time.

**Figure 2.**
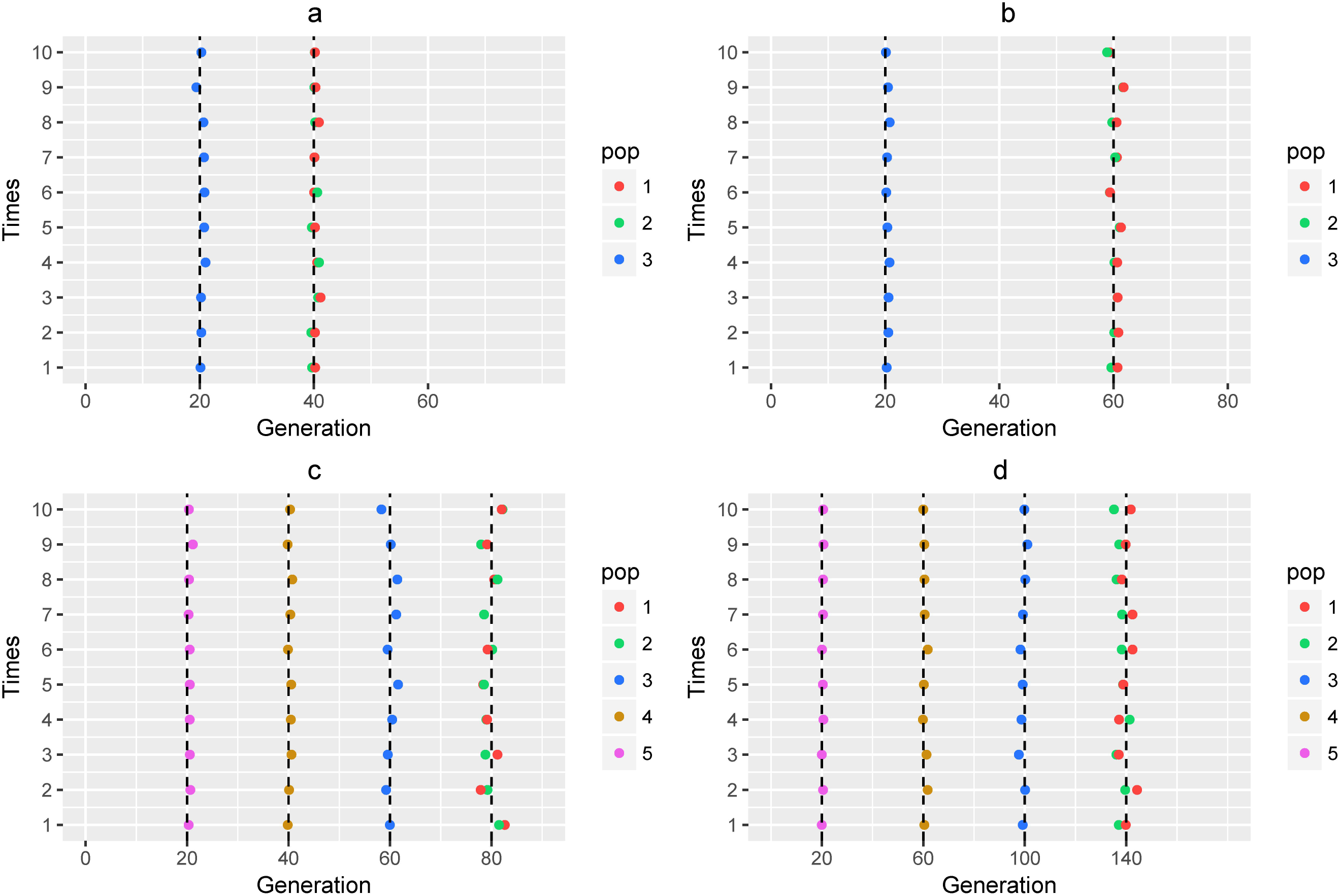
Admixture time estimated under simple model. Admixture time estimated under Scenario (I) for (a) *α*_1_ = *α*_2_ = 0.3 and *t*_2_ = 20, *T* = 40; (b) *α*_1_ = *α*_2_ = 0.3 and *t*_2_ = 40, *T* = 60. Admixture time estimated under Scenario (II) for (c) *α*_1_ = *α*_2_ = *α*_3_ = *α*_4_ = 0.3 and and *t*_2_ = 20, *t*_3_ = 40, *t*_4_ = 60, *T* = 80; (d) *α*_1_ = *α*_2_ *α*_3_ = *α*_4_ = 0.3 and *t*_2_ = 40, *t*_3_ = 80, *i*_4_ =120, *T* = 140. X-coordinate is the admixture time in generations ago, with 0 being the present time. Each case was repeated for 10 times, and *Y* = *i* means the *ith* simulation. The points in the line (*Y* = *i*) represent the admixture time estimated from the *it*h simulation, and the color of the points indicates the ancestral population. The dashed lines represent the simulated admixture time.

For the complex model, we found that our method could select the right model with high accuracy (see Fig. 3). Model selection was incorrect for only 3 simulations. In these 3 cases, the numbers of admixture waves were wrongly estimated, which led to inaccurate estimation of admixture time. Thus, selecting a correct model is of crucial importance for admixture history inference. When the admixture model could be correctly selected, only a slight overestimation occurred for the admixture time.

**Figure 3.**
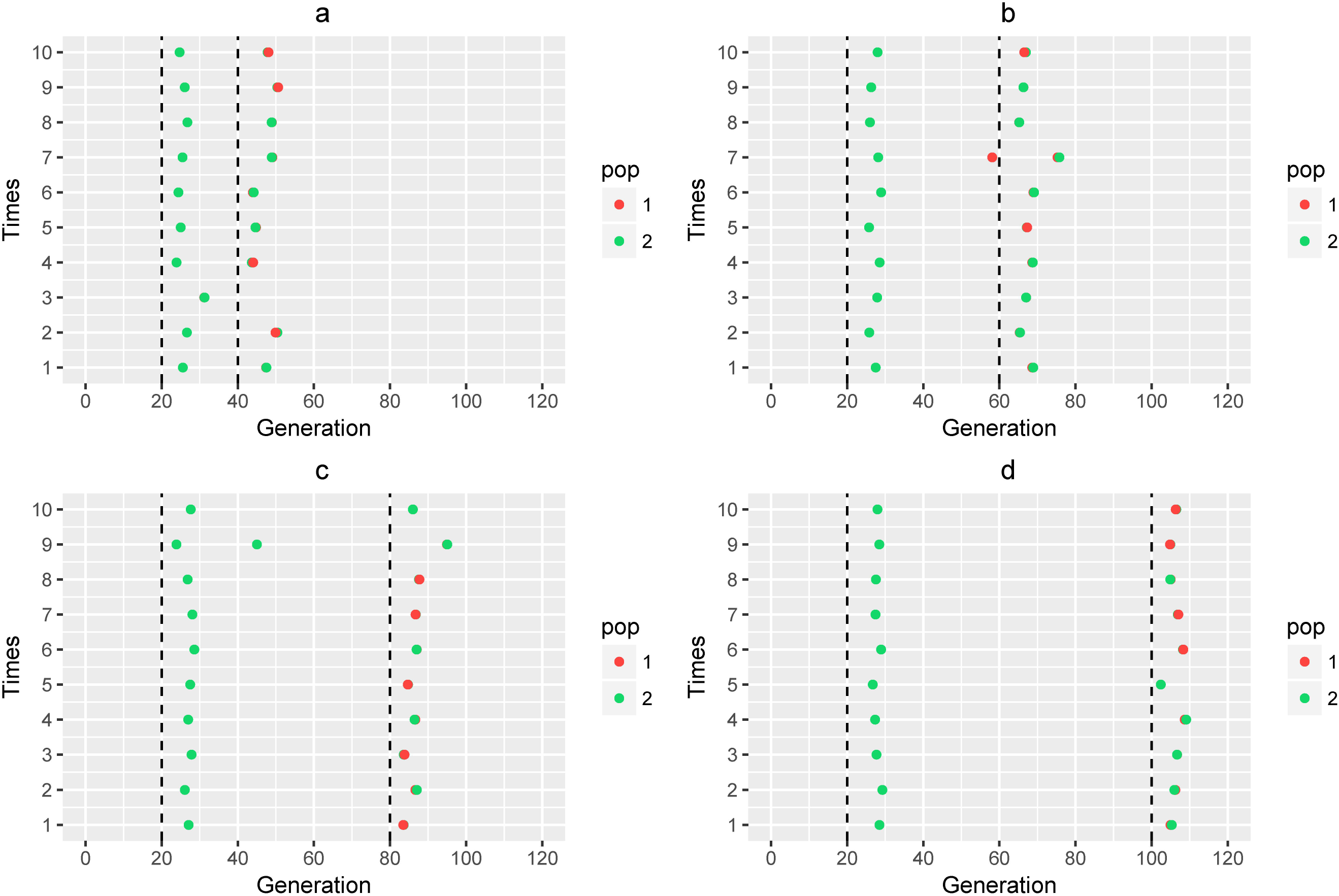
Admixture time estimated under complex model. Admixture time estimated under Scenario (III) for (a) *α*_1_ = *α*_2_ = 0.3 and *t*_2_ = 20, *T* = 40; (b) *α*_1_ = *α*_2_ = 0.3 and *t*_2_ = 40, *T* = 60; (c) *α*_1_ = *α*_2_ = 0.3 and *t*_2_ = 60, *T* = 80; and (d) *α*_1_ = *α*_2_ = 0.3 and *t*_2_ = 80, *T* = 100. X-coordinate is the admixture time ingenerations ago, with 0 being the present time. Each case was repeated for 10 times, and *Y* = *i* means the ith simulation. The points in the line (*Y* = *i*) represent the admixture time estimated from the *ith* simulation, and the color of the points indicates the ancestral population. The dashed lines represent the simulated admixture time.

We also evaluated the performance of *MultiWaver* with different admixture proportions. We found that the overestimation of admixture time in the complex model was related to the admixture proportions (see Fig. S3). When the proportions of each admixture wave became smaller, estimation error decreased. However, for the simple model, our method performed well for all situations.

In conclusion, regardless of which type of admixture model, our method performed well for model selection. Furthermore, the admixture time was estimated well for the simple model, and with only slight overestimation for the complex model.

### Robustness for Different Thresholds of Track Length

We tested the robustness of our method for different thresholds of track length. Results showed that our method was robust to thresholds for both the simple model and complex model (see Fig. 4). Due to the limitations of our method, the local ancestry inference was not so accurate for short ancestral tracks. Thus, in real data analysis, we had to discard tracks smaller than a threshold. However, short ancestral tracks contain ancient admixture information, and if the threshold was too large, lots of information would be lost. Therefore, we had to balance the trade-off between information and accuracy. In our real data analysis, we set the thresholds as 1cM.

**Figure 4.**
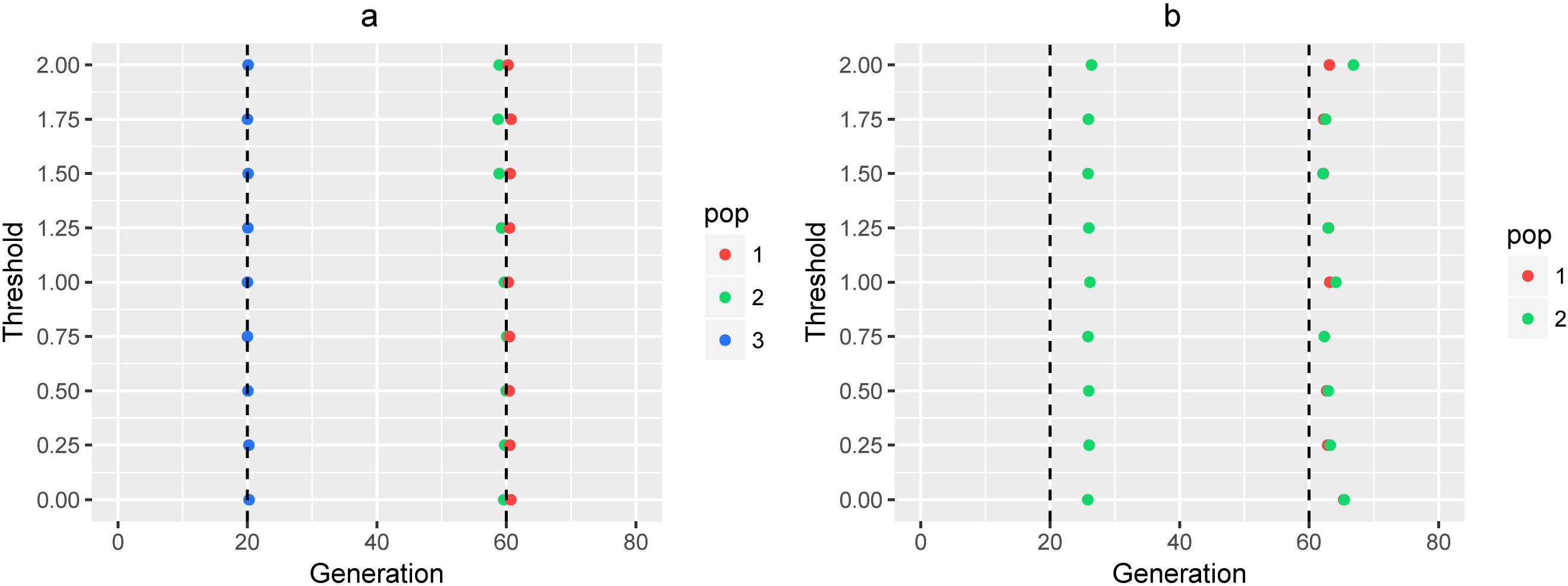
Admixture time estimated with different thresholds. (a) Admixture time estimated under Scenario (I), where *α*_1_ = *α*_2_ = 0.3 and *t*_2_ = 40, *T* = 60; (b) Admixture time estimated under Scenario (III), where *α*_1_ = *α*_2_ 0.3 and *t*_2_ = 40, *T* = 60. X-coordinate is the admixture time in generations ago, with 0 being the present time. Y-coordinate represents the thresholds, and the color of the points indicates the ancestral population. The dashed lines represent the simulated admixture time.

### Real Data Analysis

We applied our method to infer the admixture histories of some real datasets. For African Americans, HAPMIX was used to infer the ancestries with Africans (YRI) and European (CEU) as the 2 ancestral populations. The admixture model was inferred as 2 ancestral populations with a 2-wave admixtures model (see Fig. 5(a)). The African population (YRI) contributed 2 wave admixtures, and the admixture time was 11 generations ago and 7 generations ago, respectively. The time of the first admixture was about the 17th century, which was consistent with the time that most African ancestors arrived in America via slave trading. This inferred time was close to previous findings ^12,13,15,25,26^. After the slave trading, many African people settled down in America. The second admixture wave might have been caused by these people or by recent migrations from Africa to America. The admixture model inferred by our method pointed out that the admixture history of African Americans was not 1 pulse admixture, which was also reported in previous studies^12,25,26^.

**Figure 5.**
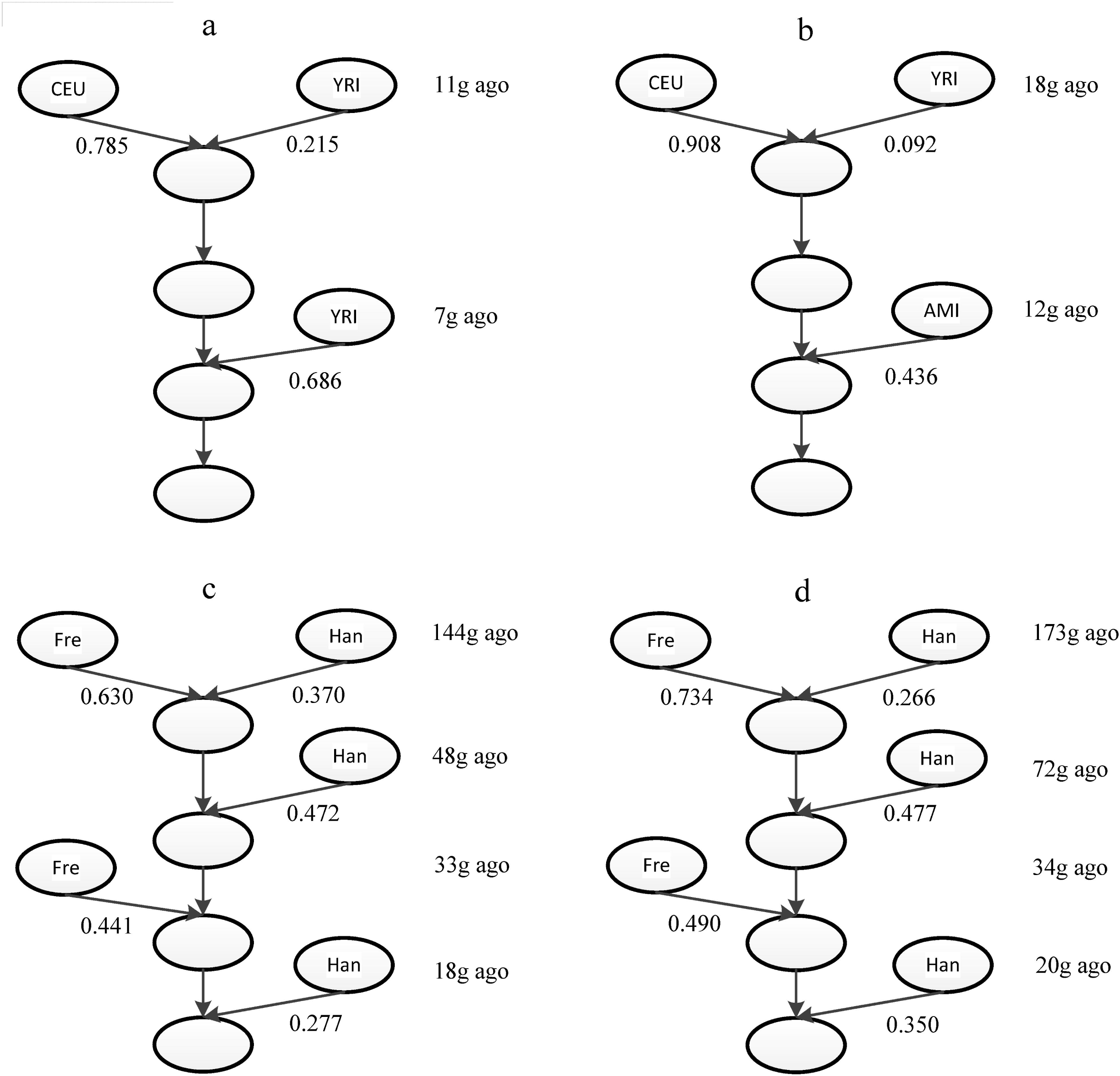
Inferred admixture history of real datasets. Inferred admixture history of (a) African Americans, (b) Mexicans, (c) Uyghurs, And (d) Hazaras. The time of the first admixture wave was the average of estimations for time *t*_0_ and *t*_1_. AMI: combined dataset of populations Maya and Pima which represent American Indian ancestry; Han: Han population, represent Asian ancestry; Fre: French population, represent European ancestry.

For Mexicans, we used PCAdmix to infer the local ancestries, and a 2-wave admixtures model was inferred (see Fig. 5(b)). Each ancestral population contributed once to the admixed population. The time of the first admixture wave was about 18 generations ago, which was close to previous findings ^25,27,31-33^. The time of the second admixture was 12 generations ago. This time period (12˜18 generation ago) was consistent with the time of the exploration of the new world. For our analysis of African Americans and Mexicans, the admixture histories inferred by our method were consistent with recorded histories, thus showing the power of our method in real data analysis.

Finally, we applied our method to reconstruct the admixture histories of Uyghurs and Hazaras (see Fig. 5(c) and (d)). Results showed that these 2 populations shared a similar admixture model, except the admixture time of Hazaras was more ancient. The earliest admixture event of Uyghurs occurred about 144 generations ago, with subsequent admixture waves from both ancestries 20-50 generations ago. While the earliest admixture event of Hazaras occurred around 173 generations ago, with following gene flows occurred 20˜70 generations ago. Compared with the results inferred by the admixture history inference method ALDER^8^, our method found an additional ancient admixture event in Uyghurs and Hazaras. To explain the discrepancies in theory, ALDER considers only the decay curve of weighted linkage disequilibrium (LD) between pairs of sites whose genetic distance were larger than 0.5 cM^8^, and thus the information of ancient signals within shorter loci pairs would be lost. Meanwhile, our method saved some of these ancient signals by deducing the conditional length distribution of ancestral tracks even if we discarded ancestral tracks shorter than 1 cM.

Conclusively, Uyghurs and Hazaras had a similar admixture history. The ancient admixture might have been caused by the migrations of Indo-Aryan speaking people into the Indian subcontinent (1500 BC). Uyghurs mainly settled in West China, and Hazaras mainly settled in Afghanistan and Pakistan. The residences of these 2 populations were all near the Silk Road, and thus we thought the recent multiple admixtures might have been caused by the trades or migrations along the Silk Road. In fact, the real history of Uyghurs and Hazaras might be more complex than inferred. However, our method could detect some effective admixtures and provide some useful information to understand the origin and development of these complex populations.

## DISCUSSION

Complex admixture history inference has long been a challenging problem in population genetics. In this work, we proposed a general discrete admixture model to describe admixture history with multiple ancestral populations and multiple-wave admixtures. We deduced that the length distribution of ancestral tracks was a mixed exponential distribution. Based on this distribution, we developed a new method, *MultiWaver,* to infer the multiple-wave admixture histories. We used LRT to select the number of admixture waves, and implemented an exhaustion method to determine the order of admixtures. When the admixture model was determined, we applied the EM-algorithm to estimate parameters. Simulations and real data analysis showed that *MultiWaver* was precise and efficient in inferring admixture history.

Comparing with previous methods, our method showed superiority in 2 aspects. Firstly, our method could be used to infer multiple-wave admixture history, while previous methods could only infer admixture history under some simple models. Secondly, no prior admixture model was required in our method, while previous methods needed to assume a special admixture model when trying to infer admixture history. Therefore, the inferred history might be biased or even unreliable if the provided model deviates from real history. However, our method avoided this problem by selecting an optimal admixture model based on ancestral tracks.

Our method introduced an elegant solution to the complex admixture history inference. However, some problems still exist. When inferring admixture history under the complex model, overestimation occurred for the admixture time. In our method, we assumed chromosome length was infinite and there was no genetic drift, and then we found that the length distribution was a mixed exponential distribution. However, Liang and Nielsen pointed out that the length distribution did not follow an exponential distribution when the admixture time was too small or too large^34^. In the complex model, the non-exponential property would be accumulated, which might be the reason behind the overestimation we observed with our method. We also found that the overestimation was related to the admixture proportion of each admixture wave. We performed simple linear regression analysis on the errors for admixture time estimations and admixture proportion (Fig. S4).

In our method, it is possible that more than 1 optimal admixture model satisfied the constraint conditions and should be recorded. However, for all simulations we conducted, this phenomenon did not appear. This was reasonable because the admixture history had a one-to-one correspondence with the length distribution of ancestral tracks. If 1 situation had more than 1 optimal model, it implied the ancestral tracks were not accurately inferred.

The efficiency of our method was also influenced by the validity of the local ancestry inference. We tested the performance of our method with the inferred ancestral tracks (see Supplementary Text 2). We found that *MultiWaver* tended to overestimate the number of waves, and thus led to overestimating the admixture time with the ancestral tracks inferred by HAPMIX (Fig. S5 (a)). For multiple-way admixtures, the inaccuracy of ancestral tracks inferred by PCAdmix led to underestimating the time of the first admixture wave (Fig. S5 (b)). It was very difficult to obtain relatively accurate ancestral tracks with a small length for all local ancestry inference methods. To improve the effectiveness of the inference, we suggest using the ancestral tracks longer than a certain threshold *C* in our method. However, when the threshold became large, ancient admixture information would be lost rapidly. With the development of sequencing technology and computational methods, short ancestral tracks could be precisely detected in the near future. Then, our method would be promising in recovering even more ancient admixture history, such as the admixture between modern humans and ancient humans^35,36^.

## ACKNOWLEDGEMENTS

This work was supported by National Natural Science Foundation of China (NSFC) grants (91331204 and 11426237), the Strategic Priority Research Program (XDB13040100) and Key Research Program of Frontier Sciences (QYZDJ-SSW-SYS009) of the Chinese Academy of Sciences (CAS), the National Science Fund for Distinguished Young Scholars (31525014), the Program of Shanghai Academic Research Leader (16XD1404700); 973 Project (2011CB808000), the Fundamental Research Funds for the Central Universities (2011JBZ019, 2014RC008 and 2015IBM099), the National Excellent Doctoral Dissertation Foundation of PR China (201213), National Center for Mathematics and Interdisciplinary Sciences of CAS, and the Key Laboratory of Random Complex Structures and Data Science, CAS (2008DP173182). S.X. also gratefully acknowledges the support of the National Program for Top-notch Young Innovative Talents of The “*Wanren Jihua*” Project. We thank LetPub (www.letpub.com) for its linguistic assistance during the preparation of this manuscript.

